# The oncogenic E3 ligase TRIP12 suppresses epithelial-mesenchymal transition (EMT) and metastasis-related processes through ZEB1/2

**DOI:** 10.1101/2020.11.04.368209

**Authors:** Kwok Kin Lee, Deepa Rajagopalan, Shreshtha Sailesh Bhatia, Wee Joo Chng, Sudhakar Jha

## Abstract

Thyroid hormone receptor interactor 12 (TRIP12) is an E3 ligase most notably involved in the proteolytic degradation of the tumor suppressor p14ARF. Through this process, it is proposed that TRIP12 plays an oncogenic role in tumor initiation and growth. However, its role in other cancer processes such as metastasis is unknown. In this study, using publicly available cancer patient datasets, we found TRIP12 to be associated with distant metastasis-free survival in breast cancer, suggesting a contrary cancer process inhibitory role in metastasis. Following TRIP12 depletion in the MCF10A breast epithelial cell model, an epithelial-mesenchymal transition (EMT) shift occurred with concomitant changes in EMT cell adhesion markers identified through RNA-seq. In line with EMT changes, TRIP12-depleted cells lose polarity and dislodge from bulk cells at a higher frequency. Furthermore, ectopic TRIP12 expression sensitized cells to anoikis, a major barrier against metastasis. Mechanistically, TRIP12 suppresses EMT through inhibiting ZEB1/2 gene expression and ZEB1/2 depletion rescues EMT markers and cellular behavior. Overall, our study delineates TRIP12’s role in inhibition of EMT and metastasis-related processes, and implies a suppression role in breast cancer metastasis.

## INTRODUCTION

Metastasis is a multi-step process resulting in the dissemination of cancer cells from the primary tumor site to a secondary site (Thiery, 2002). Within the metastatic cascade, epithelial-mesenchymal transition (EMT), an embryonic transcriptional program hijacked by cancer cells, plays a crucial first step to initiate metastasis (Thiery, 2002). During EMT, epithelial cells undergo cellular reprogramming to transit into different quasi-mesenchymal cell states within an EMT spectrum (Dongre and Weinberg, 2019; Nieto et al., 2016; Tan et al., 2014; Thiery, 2002; Thiery et al., 2009). Neoplastic cells in the quasi-mesenchymal cell states have remodeled cytoskeleton, increased motility and exhibit high aggressiveness (Dongre and Weinberg, 2019; Nieto et al., 2016). Particularly in breast cancer, metastasis remains the highest cause of death with a 5-year survival rate of only 27% for breast cancer patients with distant metastasis, as compared to patients with localized or regional spread whose 5-year survival rates are 99% and 85% respectively (Siegel et al., 2020), highlighting that metastasis remains an incompletely understood disease (Lambert et al., 2017).

Thyroid hormone receptor interactor 12 (TRIP12) is a homologous to E6AP C terminus (HECT) domain E3 ubiquitin ligase essential in various cellular processes and pathways. It is required for embryonic development, as mouse embryos with TRIP12 homozygous mutation are embryonic lethal and have delayed development at E8.5 (Kajiro et al., 2011). Additionally, TRIP12 maintains pancreatic cell homeostasis through regulating the protein stability of pancreas transcription factor 1a (PTF1A) (Hanoun et al., 2014). In the DNA damage context, TRIP12 prevents the uncontrolled spreading of chromatin ubiquitination through its degradation of ring finger protein 168 (RNF168), the E3 ligase responsible for histone ubiquitination upon DNA damage and subsequent recruitment of 53BP1 (Gudjonsson et al., 2012). In human papillomavirus induced head and neck cancers where p16 levels are elevated, negative regulation of TRIP12 level by p16 increases sensitivity to radiotherapy due to the loss of DNA repair pathway (Wang et al., 2017). More recently, a study discovered that TRIP12 is a poly(ADP-ribose) (PAR)-dependent E3 ubiquitin ligase responsible for degradation of poly(ADP-ribose) polymerase 1 (PARP1), much like the pioneer PAR-dependent E3 ligase ring finger protein 146 (RNF146/IDUNA) (Callow et al., 2011; Kang et al., 2011; Zhang et al., 2011), thus constraining PARP inhibitor efficiency (Gatti et al., 2020). Apart from the above-mentioned pathways, TRIP12 also serves as the E3 ligase for the ubiquitin fusion degradation (UFD) pathway (Park et al., 2009) which is responsible for the degradation of substrates with an N-terminal linked ubiquitin (Johnson et al., 1995) and has been suggested to be the core degradation pathway for lysine-less substrates such as p14ARF (Collado and Serrano, 2010). In addition, 2 studies show that TRIP12 is important for the maintenance of protein complex stoichiometry namely, the SWI/SNF complex and the APP-BP1 neddylation complex through the degradation of their unbound subunits BAF57 and APP-BP1 respectively (Keppler and Archer, 2010; Park et al., 2008).

In the cancer context, TRIP12 is a key regulatory player in oncogene-induced senescence (OIS), a major barrier against cancer initiation. In normal human fibroblast cells, the steady-state level of p14ARF is tightly controlled through protein degradation mediated by TRIP12 (Chen et al., 2010a). In oncogene-induced scenarios such as c-Myc amplification, this TRIP12-p14ARF regulation is disrupted resulting in an increase of p14ARF level, thereby leading to OIS and the prevention of carcinogenesis (Chen et al., 2013; Chen et al., 2010a). In addition, several p14ARF interacting partners such as nucleophosmin (NPM), tumor necrosis factor receptor associated death domain (TRADD), N-Myc and STATs interactor (NMI), and nucleostemin have also been reported to suppress cell growth or tumor formation through the stabilization of p14ARF by disrupting TRIP12-p14ARF degradation in different cell types including, normal fibroblast, acute myeloid leukemia, lung cancer cell lines and in a mouse cancer model (Chen et al., 2013; Chen et al., 2010a; Chen et al., 2010b; Chio et al., 2012; Li et al., 2012; Lo et al., 2015). More recently, studies in liver cancer found that the deubiquitinating enzyme USP7 stabilizes TRIP12, which leads to constitutive p14ARF ubiquitination and degradation, thereby promoting the growth of liver cancer cells (Cai et al., 2015). Interestingly, another study found that TRIP12 could degrade USP7 (Liu et al., 2016), suggesting a possible regulatory loop between TRIP12 and USP7. TRIP12 has also been found to have the highest frequency of mutations in lung adenocarcinoma patients, while frameshift mutations of TRIP12 are present in colorectal and gastric cancers with microsatellite instability (Li et al., 2014; Yoo et al., 2011). Taken together, there is a consensus in the possible tumor-promoting role of TRIP12 through its proteolytic effect on p14ARF.

Although the role of TRIP12 as a potential promoter of cancer initiation has been studied, it's role in different cancer types and in cancer progression remains unknown. In this present study, we identified an inhibitory role for TRIP12 in suppressing epithelial-mesenchymal transition (EMT) and metastasis-related processes through ZEB1/2 gene expression, contrary to its tumor promoting role in cancer initiation.

## RESULTS

### TRIP12 expression in breast cancer patients correlates with distant metastasis-free survival

As a preliminary step to study the role of TRIP12 in different cancers, online bioinformatics tools for meta-analysis were used to determine the clinical relevance of TRIP12 expression. Among different clinical parameters and different cancers, we observe that TRIP12 is associated with distant metastasis-free survival of breast cancer patients (Fig. 1). Through the PrognoScan tool (Mizuno et al., 2009), TRIP12 is found to correlate with distant metastasis-free survival in 3 publicly available breast cancer datasets. Distant metastasis-free survival is significantly lower (COX p-value=0.01819; HR = 0.20(95% CI: 0.05-0.76)) in breast cancer patients with low TRIP12 as compared to high TRIP12 levels in the GSE11121 study (Fig. 1A). Similarly, in GSE6532-GPL570, there is a significantly lower (COX p-value=0.03578; HR=0.27(95% CI: 0.08-0.92)) probability of distant metastasis in high TRIP12 breast tumors as compared to low TRIP12 tumors (Fig. 1B). Lastly, distant metastasis-free survival is lower (p-value = 0.03563; COX p-value = 0.26621; HR=0.70(95% CI: 0.37-1.32)) in TRIP12 low tumors than in TRIP12 high tumors in the GSE7390 breast cancer study (Fig. 1C). A similar correlation is observed in the GSE5327 breast cancer dataset, obtained through the PROGgeneV2 online database (Goswami and Nakshatri, 2014), which shows that lung metastasis-free survival is lower (p-value=0.00337; HR=0.21(95% confidence interval (CI): 0.07-0.60)) in tumors with low TRIP12 as compared to tumors with high TRIP12 levels (Fig. 1D). These data suggest that TRIP12 could have an inhibitory role in breast cancer distant metastasis.

**Fig. 1.**
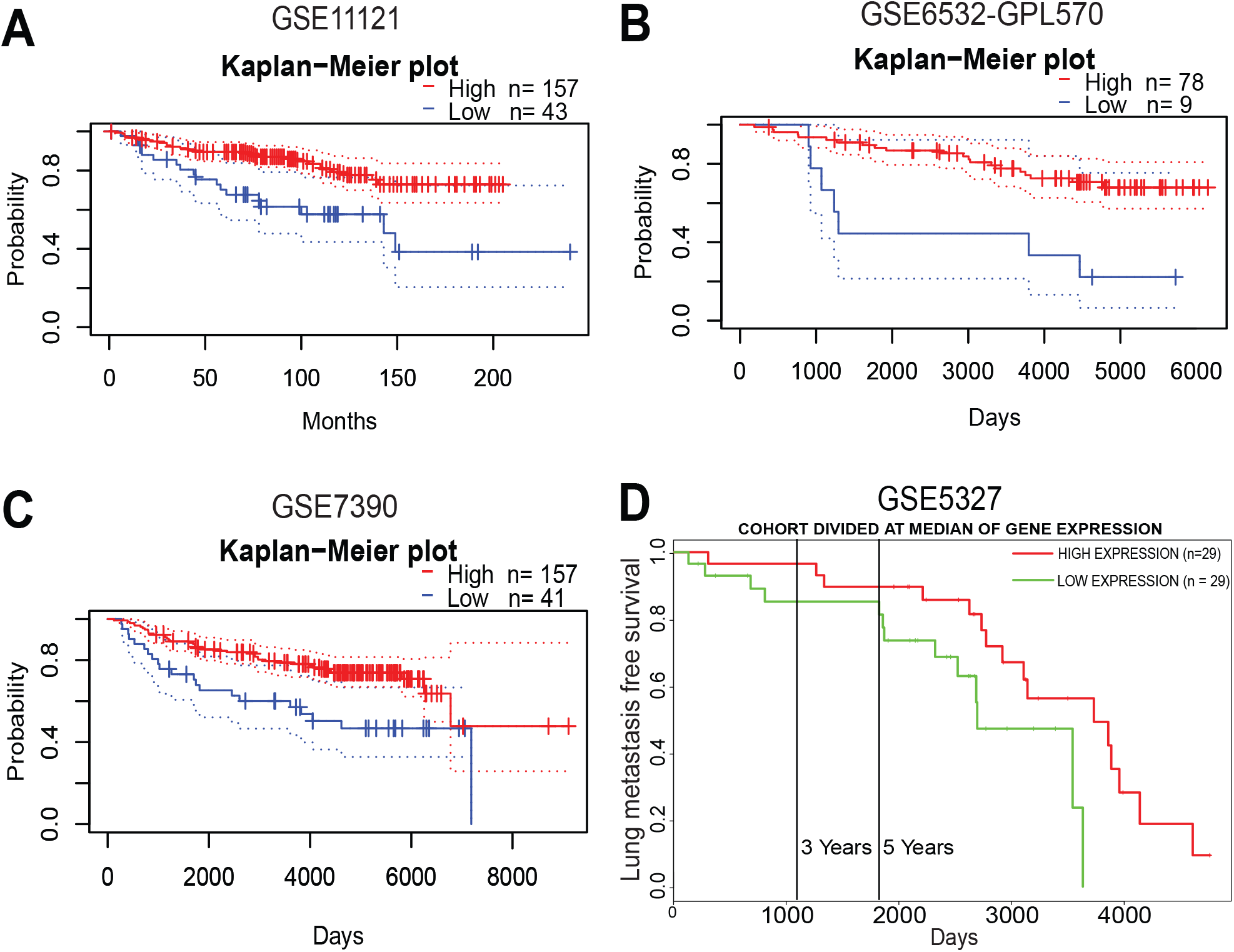
*TRIP12* expression in breast cancer patients correlates with distant metastasis-free survival. (A) Kaplan-Meier distant metastasis-free survival curve for breast cancer patients in GSE11121. (B) Kaplan-Meier distant metastasis-free survival curve for breast cancer patients in GSE6532-GPL570. (C) Kaplan-Meier distant metastasis-free survival curve for breast cancer patients in GSE7390. (D) Kaplan-Meier lung metastasis-free survival curve for breast cancer patients in GSE5327.

### RNA-seq reveals cell adhesion molecules as the most significant pathway regulated by TRIP12

To study the role of TRIP12 in the breast metastasis context, the breast epithelial MCF10A cell model, which has been used to investigate the EMT process (Maeda et al., 2005; Sarrió et al., 2008) is utilized. Two stable cell lines expressing different shRNAs targeting TRIP12 were generated. The stable cell lines robustly show a decrease in TRIP12 expression both in the RNA (Fig. 2A) and protein (Fig. 2B) levels. EMT is a form of cellular reprograming characterized by wide-spread transcriptional changes in the gene expression of multiple EMT markers (Lamouille et al., 2014). In order to characterize global gene expression changes regulated by TRIP12, RNA-sequencing (RNA-seq) was performed on the generated stable cells depleted of TRIP12. Genes that are significantly differentially expressed (FDR < 0.05 and |log2 (fold change) | > 1) in the 2 stable cell lines with different shRNA targeting TRIP12 are represented by the volcano plots in Fig. 2C. The list of all differentially expressed gene from the 2 stable cell lines are shown in Table S1. Overlap of the significantly differentially regulated genes between the 2 stable cell lines identified 197 downregulated genes and 71 upregulated genes (Fig. 2C) regulated by TRIP12.

**Fig. 2.**
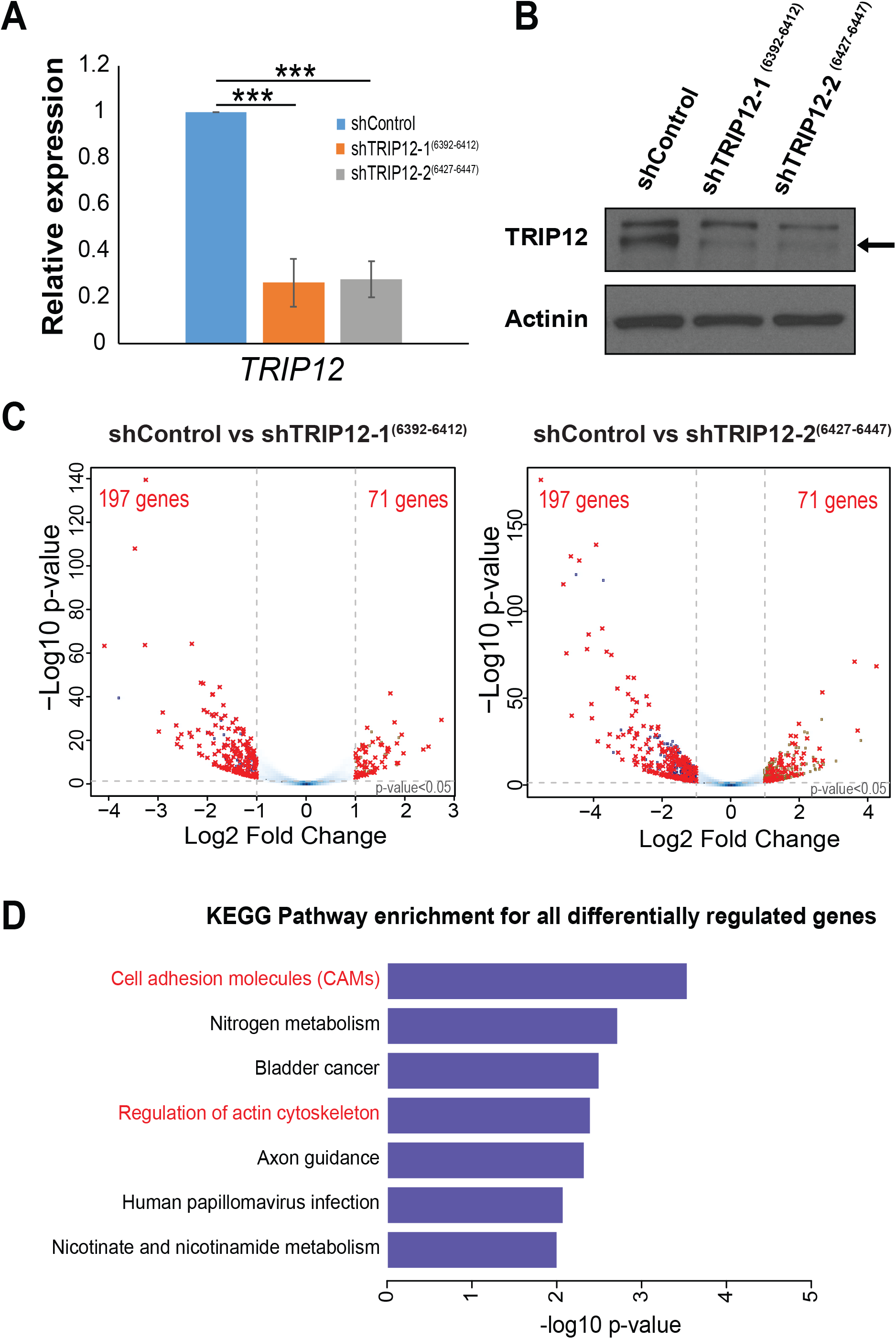
RNA-seq reveals cell adhesion molecules as the most significant pathway regulated by TRIP12. (A) Relative expression of *TRIP12* mRNA in MCF10A cells stably depleted of TRIP12. Relative expression levels were normalized to actin mRNA levels and data quantified relative to shControl. (N = 3). Data represents means ± SD. ***, p-value < 0.001. (B) TRIP12 and actinin protein levels in MCF10A cells stably depleted of TRIP12. Actinin serves as a loading control. Arrow indicates the TRIP12 band. (N = 3). (C) Volcano plots showing the differentially regulated genes upon TRIP12 depletion in MCF10A for shTRIP12-1^(6392-6412)^ (left panel) and shTRIP12-2^(6427-6447)^ (right panel), as compared to shControl. Differentially regulated genes were identified using the criteria: FDR < 0.05 and |Log2 fold change | > 1. Dots marked in red represent the overlap of differentially regulated genes between shTRIP12-1^(6392-6412)^ and shTRIP12-2^(6427-6447)^. (N = 2 for each shRNA). (D) KEGG pathway analysis for all differentially regulated genes which overlap between shTRIP12-1^(6392-6412)^ and shTRIP12-2^(6427-6447)^ shown in C. Pathways related to EMT are highlighted in red.

Pathway analysis of the overlapped differentially regulated genes enriches for several pathways, including “cell adhesion molecules (CAMs),” “nitrogen metabolism,” “bladder cancer,” “regulation of actin cytoskeleton,” “axon guidance,” “human papillomavirus infection,” and “nicotinate and nicotinamide metabolism” (Fig. 2D). Among the pathways regulated, “cell adhesion molecules (CAMs)” and “regulation of actin cytoskeleton” represent EMT-related processes, with “cell adhesion molecules (CAMs)” as the most significantly enriched pathway (Fig. 2D). Cell adhesion molecules play an important role in maintaining cell-cell contact and are frequently altered during epithelial-mesenchymal transition (EMT) (Dongre and Weinberg, 2019; Huang et al., 2012; Nieto et al., 2016; Thiery, 2002; Thiery et al., 2009). Examples include the epithelial marker E-cadherin (*CDH1*) and mesenchymal marker N-cadherin (*CDH2*), which are canonical EMT markers. Since TRIP12 affects the expression of cell adhesion molecules, we follow up to study the alteration in cell adhesion molecules expression by TRIP12 and its effects.

### TRIP12 regulates EMT cell adhesion genes involved in adherens junctions, tight junctions, desmosomes, cytoskeleton, and extracellular matrix interaction

Corroborating with changes in cell adhesion molecules gene expression, morphological changes are observed in MCF10A cells depleted of TRIP12 (Fig. 3A). Cells lose their cobblestone morphology, are more elongated, and exhibit a “fibroblast-like” spindle morphology (Fig. 3A), typical of cells that have undergone EMT. Images were taken at a high density because MCF10A cell density has been reported to affect morphology, whereby cells display an epithelial morphology at high density while adopting a mesenchymal morphology at low density (Maeda et al., 2005; Sarrió et al., 2008). TRIP12-depleted cells at high density still show a mesenchymal morphology (Fig. 3A), confirming that these morphological changes are not due to cell density.

**Fig. 3.**
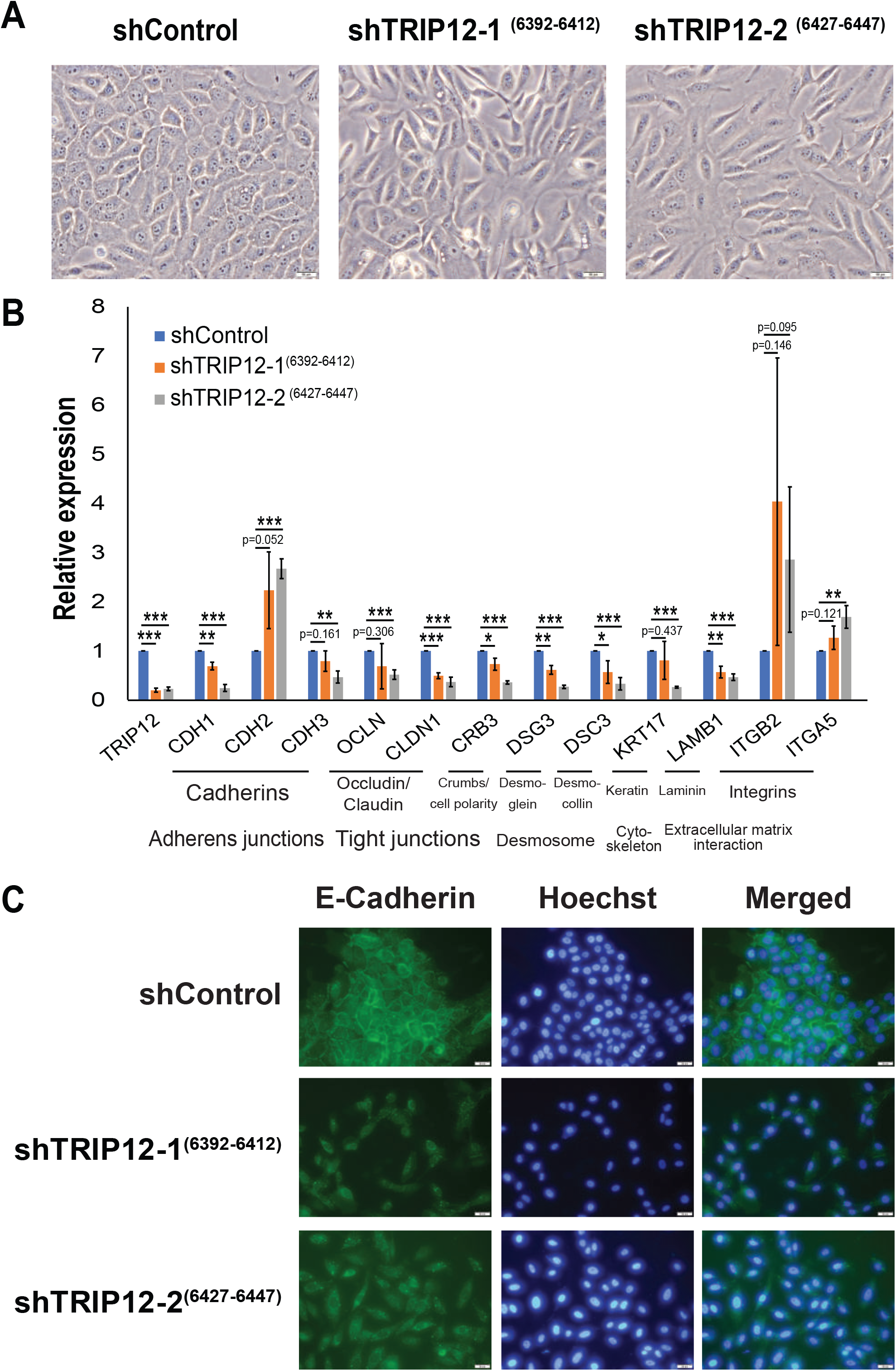
TRIP12 regulates EMT cell adhesion genes involved in adherens junctions, tight junctions, desmosomes, cytoskeleton, and extracellular matrix interaction. (A) Bright-field images showing the cellular morphology of MCF10A shControl, shTRIP12-1^(6392-6412)^ and shTRIP12-2^(6427-6447)^. Images were taken at 20X magnification. The scale bar at bottom right corner = 50μm. (N = 1). (B) Relative expression of the mRNA levels of EMT markers in MCF10A cells stably depleted of TRIP12. Relative expression levels were normalized to actin mRNA levels and data quantified relative to shControl. (N = 3). Data represents means ± SD. *, p-value < 0.05; **, p-value < 0.01; ***, p-value < 0.001. (C) Fluorescent images of MCF10A shControl, shTRIP12-1^(6392-6412)^ and shTRIP12-2^(6427-6447)^ immunostained using an E-cadherin antibody and with Hoechst dye. Images were taken at 20X magnification. The scale bar at bottom right corner = 50μm. (N = 1).

Next, quantitative-Polymerase Chain Reaction (qPCR) was utilized to validate genes involved in cell adhesion. TRIP12 downregulation alters the expression of different cellular junction genes (Fig. 3B). Notably, changes in adherens junctions include the loss of epithelial markers E-cadherin (*CDH1*) and P-cadherin (*CDH3*), and the gain of mesenchymal marker N-cadherin (*CDH2*) (Fig. 3B). This is typically termed “cadherin switching” and is observed in EMT during embryonic development (Wheelock et al., 2008). In addition, TRIP12-depleted cells show a decrease in membrane E-cadherin protein (Fig. 3C), corroborating with changes in E-cadherin gene expression. Tight junction structural proteins occludin (*OCLN*) and claudin 1 (*CLDN1*), and tight junction-associated protein crumbs cell polarity complex component 3 (*CRB3*) are also downregulated upon TRIP12 depletion (Fig. 3B). Desmosomes are a component of the epithelial junctional complex and aid in cell-cell adhesion (Zihni et al., 2016). Structural components of the desmosome, desmocollin-3 (*DSC3*) and desmoglein-3 (*DSG3*) show a decrease upon TRIP12 depletion (Fig. 3B). Other important signaling and EMT-related molecules such as keratin (*KRT17*), laminin (*LAMB1*) and integrins (*ITGB2* and *ITGA5*) are also altered upon TRIP12 depletion (Fig. 3B) These results validate the RNA-seq data in which TRIP12 depletion leads to major gene expression changes in cell adhesion molecules.

### Alterations in TRIP12 level results in loss of cellular polarity, increase in single-cell dislodgment, and sensitivity to anoikis

To study if the expression changes in cell adhesion molecules are specific to TRIP12, stable cell lines with ectopic expression of TRIP12 are created. The cell lines with ectopic TRIP12 expression show an increase in TRIP12 level which can be differentiated from endogenous TRIP12 using qPCR primers either targeting TRIP12 3`UTR or CDS (Fig. 4A; Fig. S1). Upon ectopic TRIP12 expression, the majority of cell adhesion genes expressions are partially rescued (Fig. 4A). Epithelial markers *CDH1*, *CDH3*, *OCLN*, *CLDN1*, *CRB3*, *DSG3*, *DSC3*, and *KRT17*, show a higher level in cells with TRIP12 depletion and ectopic LHCX-TRIP12 expression, as compared to cells with TRIP12 depletion and ectopic LHCX-vector expression (Fig. 4A). Among these, the level of *OCLN*, *CDH3* (for shTRIP12-2^(6427-6447)^ LHCX-TRIP12) and *CRB3* (for shTRIP12-2^(6427-6447)^ LHCX-TRIP12), are significantly rescued.

**Fig. 4.**
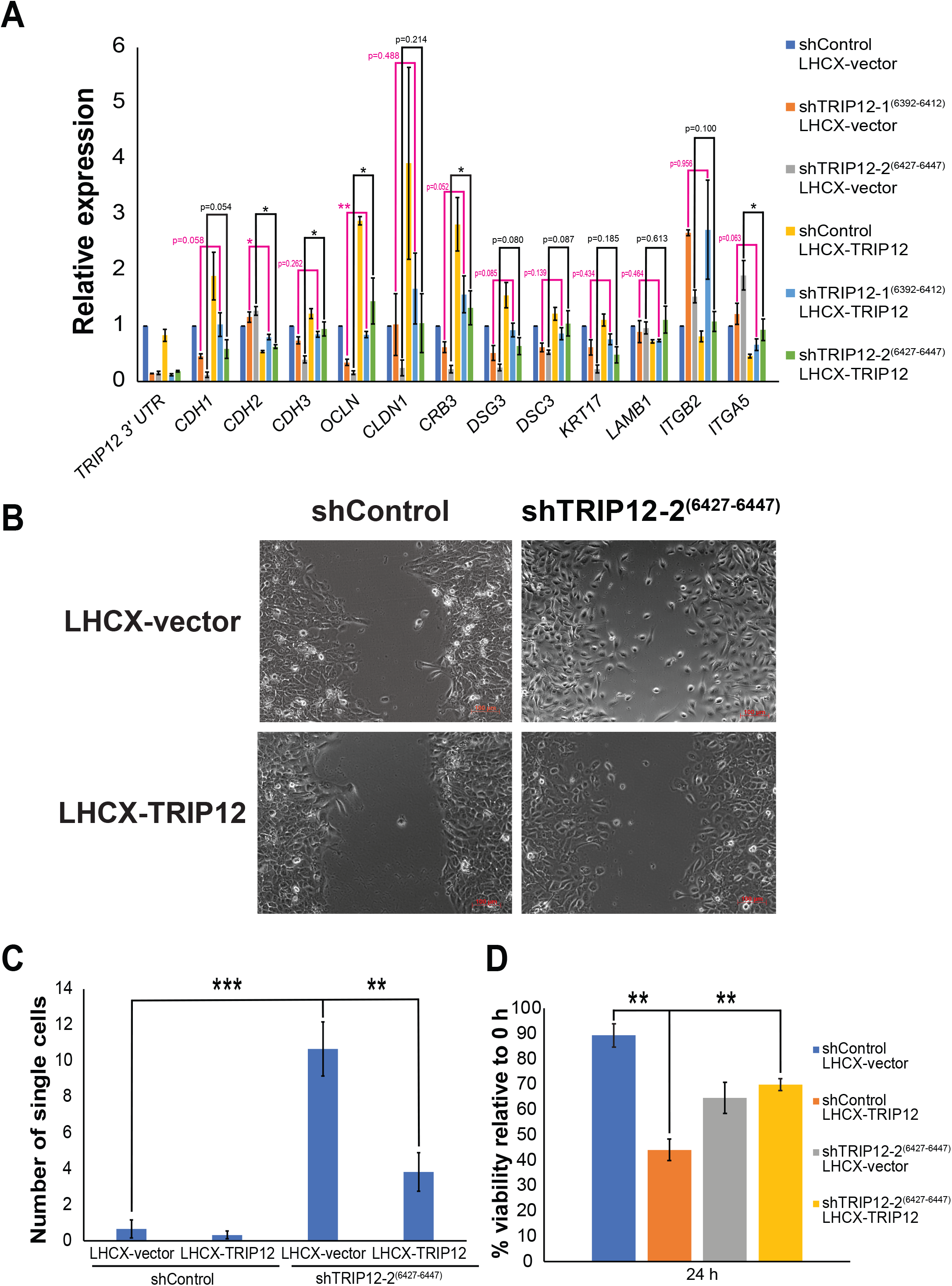
Alterations in TRIP12 level results in loss of cellular polarity, increase in single-cell dislodgment, and sensitivity to anoikis. (A) Relative expression of the mRNA levels of EMT markers in MCF10A cells stably depleted of TRIP12 and with LHCX-vector expression or LHCX-TRIP12 expression. Relative expression levels were normalized to actin mRNA levels and data quantified relative to shControl LHCX vector. (N = 3). Data represents means ± SEM. *, p-value < 0.05; **, p-value < 0.01. (B) Representative snapshot images of cell migration/ wound-healing assay for MCF10A shControl and shTRIP12-2^(6427-6447)^, with either LHCX-vector or LHCX-TRIP12 expression, when remaining wound area is 35% −45% of total area. Images were taken at 10X magnification. The scale bar at bottom right corner = 100μm. (C) Quantification of the number of single cells in the remaining wound area from images when the remaining wound area is 35% −55% of the total area. (N = 6). Data represents means ± SEM. **, p-value < 0.01; ***, p-value < 0.001. (D) Percent cell viability data for MCF10A shControl and shTRIP12-2^(6427-6447)^, with either LHCX-vector or LHCX-TRIP12 expression seeded on agarose coated wells for 24 h. Cell viability was measured using a MTS assay and expressed as a percentage to 0 h. Data represents means ± SEM. (N = 3).

Mesenchymal markers *CDH2*, *ITGB2* (for shTRIP12-2^(6427-6447)^ LHCX-TRIP12) and *ITGA5*, show a lower level in cells with TRIP12 depletion and ectopic LHCX-TRIP12 expression, as compared to cells with TRIP12 depletion and LHCX-vector expression (Fig. 4A). Specifically, mesenchymal markers *CDH2* and *ITGA5* (for shTRIP12-2^(6427-6447)^ LHCX TRIP12) show a significant rescue in level. Notably, ectopic expression of TRIP12 in shControl cells also results in an opposite trend to TRIP12 depletion for both epithelial and mesenchymal markers (Fig. 4A), affirming the role of TRIP12 in regulating cell adhesion genes, and depicting the plasticity of MCF10A to transverse along the EMT spectrum.

To investigate if TRIP12 has a role to play in metastasis processes, a wound-healing assay was performed in MCF10A shTRIP12-2^(6427-6447)^ cells, which show a larger extent of change in EMT markers expression as compared to MCF10A shTRIP12-1^(6392-6412)^ cells (Fig. 3A). Cells with loss of TRIP12 (shTRIP12-2^(6427-6447)^ LHCX-vector) dislodge from bulk cells towards the wound area as single cells at a significantly higher frequency than control cells (shControl LHCX-vector) (Fig. 4B and 4C; Movies S1 and S2). In addition, TRIP12-depleted cells also lose polarity and move in a random orientation towards the wound area as compared to control cells (Movies S1 and S2). Importantly, ectopic expression of TRIP12 rescues this cellular behavior (Fig. 4B and 4C; Movies S1-S4). Significantly fewer MCF10A shTRIP12-2^(6427-6447)^ LHCX-TRIP12 cells dislodge as single cells (Fig. 4C). MCF10A shTRIP12-2^(6427-6447)^ LHCX-TRIP12 cells are more oriented and move towards closing the wound in a sheet-like fashion similar to shControl cells (Movies S1-S4). An important step in the metastatic cascade is the dissemination of single cells or collective migration into lymphatic or blood circulation (Lambert et al., 2017). The observation that loss of TRIP12 results in more single-cell dislodgment which is rescued by ectopic TRIP12 (Fig. 4B and 4C), suggests a prohibitory role for TRIP12 in the initial stage of the metastatic cascade.

Anoikis is the induction of programmed cell death in the absence of cell attachment or attachment to inappropriate surfaces (Paoli et al., 2013). Resistance to anoikis is a feature acquired by metastatic cells to survive in circulation and for the successful colonization on distant organs (Paoli et al., 2013). Expression of *OCLN*, one of the cell adhesion genes regulated by TRIP12 (Fig. 4A), has been shown to decrease cancer cell’s resistance to anoikis (Osanai et al., 2006). Similarly, ectopic expression of TRIP12 which increases *OCLN* expression (Fig. 4A), results in a decrease in viability when cells are seeded on agarose-coated wells to induce anoikis (Fig. 4D). Subsequently, decreasing the total TRIP12 level using shRNA results in a rescue in viability (Fig. 4D), with corresponding *OCLN* expression level changes (Fig. 4A), indicating TRIP12’s specificity in anoikis. Additionally, this effect is specific to anoikis and not a case of general cell death due to TRIP12 ectopic expression, as supported by Fig. S2 where there is no difference in cell viability when cells are seeded on tissue-culture plates and viability measured. These results depict TRIP12’s role in anoikis induction (Fig. 4D) and suggest an inhibitory role for TRIP12 to prevent metastatic cells from gaining anoikis resistance in the metastatic cascade.

### TRIP12 regulates EMT and metastasis-related processes through ZEB1/2 gene expression

The canonical EMT transcription factors (EMT-TFs), comprising of the snail family transcription repressors (SNAI1, SNAI2), the zinc finger E-box binding homeobox family (ZEB1, ZEB2) and the twist family bHLH transcription factor family (TWIST1), represent the molecular drivers of EMT and of metastasis (Thiery et al., 2009; Yang and Weinberg, 2008). These transcription factors drive EMT mainly through their direct or indirect effects on different cellular junction proteins, such as the repression of *CDH1* and polarity factor *CRB3* (Batlle et al., 2000; Cano et al., 2000; Comijn et al., 2001; Eger et al., 2005; Hajra et al., 2002; Thiery et al., 2009; Vandewalle et al., 2005; Yang et al., 2004; Yang and Weinberg, 2008). Among the EMT-TFs, RNA-seq data shows *ZEB1* and *ZEB2* gene expression significantly upregulated, with *ZEB2* expression increasing more than 2-folds upon TRIP12 depletion (Table S1). To validate, we performed gene expression analysis for EMT-TFs using qPCR. TWIST2 is also included as it has also been shown to promote EMT (Fang et al., 2011). Following TRIP12 depletion, only *ZEB1* and *ZEB2* expression increases consistently in both shTRIP12-1^(6392-6412)^ and shTRIP12-2^(6427-6447)^ (Fig. 5A). No significant change was noted for *TWIST1* and *TWIST2* (Fig. 5A). On the contrary, *SNAI1* and *SNAI2* changes in opposing directions between shTRIP12-1^(6392-6412)^ and shTRIP12-2^(6427-6447)^ (Fig. 5A). This could be due to differences in quasi-mesenchymal states within the spectrum which these 2 different shTRIP12 stable lines are in, as different combinations and timely contribution of different EMT-TFs is known to determine the eventual EMT state (Nieto et al., 2016; Stemmler et al., 2019). This is further supported by the degree of change of the EMT markers between shTRIP12-1^(6392-6412)^ and shTRIP12-2^(6427-6447)^, which indicates that shTRIP12-2^(6427-6447)^ is in a further mesenchymal state in the EMT spectrum as compared to shTRIP12-1^(6392-6412)^ (Fig. 3B). Additionally, the degree of change in EMT markers (Fig. 3B) correlates with the degree of change of ZEB1 and ZEB2 in shTRIP12-1^(6392-6412)^ and shTRIP12-2^(6427-6447)^ (Fig. 4A), suggesting that ZEB1/2 expression regulation by TRIP12 (Fig. 5A) regulates TRIP12-dependent EMT inhibition.

**Fig. 5.**
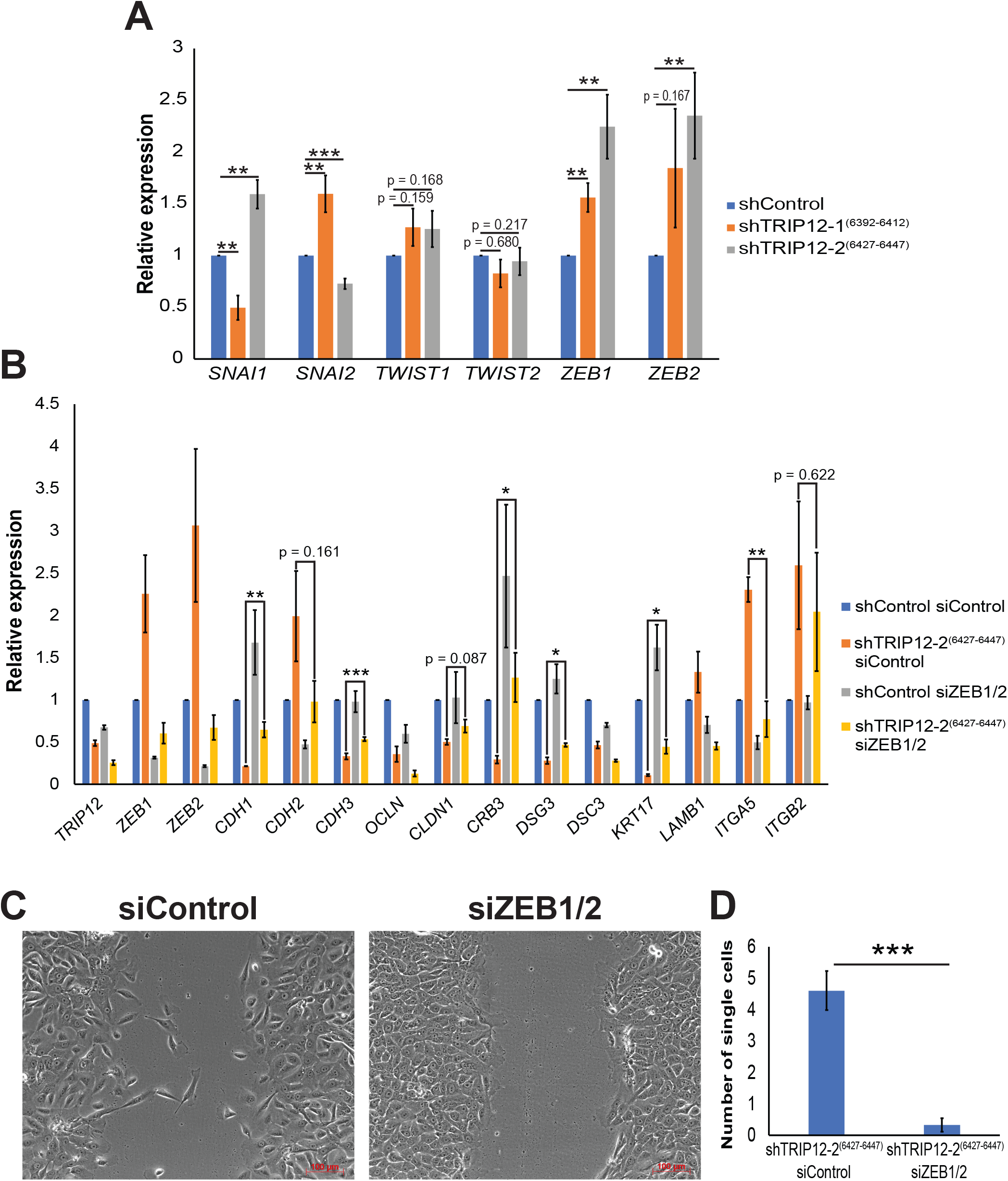
TRIP12 inhibits ZEB1/2 gene expression to regulate EMT and cellular behavior. (A) Relative expression of the mRNA levels of EMT transcription factors in MCF10A cells stably depleted of TRIP12. Relative expression levels were normalized to actin mRNA levels and data quantified relative to shControl. (N = 7). Data represents means ± SEM. **, p-value < 0.01; ***, p-value < 0.001. (B) Relative expression of the mRNA levels of EMT markers in MCF10A cells stably depleted of TRIP12 and with ZEB1/2 depletion. Relative expression levels were normalized to actin mRNA levels and data quantified relative to shControl siControl. (N = 3). Data represents means ± SEM. *, p-value < 0.05; **, p-value < 0.01; ***, p-value < 0.001. (C) Representative snapshot images of cell migration/ wound-healing assay for MCF10A shTRIP12-2^(6427-6447)^, with either siControl or siZEB1/2 when remaining wound area is 35% −45% of total area. Images were taken at 10X magnification. The scale bar at bottom right corner = 100μm. (D) Quantification of the number of single cells in the remaining wound area from images when the remaining wound area is 35% - 45% of the total area. (N = 5). Data represents means ± SEM. ***, p-value < 0.001.

To study the role of ZEB1 and ZEB2 in TRIP12-dependent EMT inhibition, a co-depletion of *ZEB1* and *ZEB2* using siRNA was performed. Starkly, depletion of both *ZEB1* and *ZEB2* rescued the expression of epithelial genes *CDH1*, *CDH3*, *CLDN1*, *CRB3*, *DSG3*, and *KRT17* (Fig. 5B). Among these, *CDH1*, *CDH3*, *CRB3*, *DSG3*, and *KRT17* gene expression are significantly rescued (Fig. 5B). On the other hand, mesenchymal genes *CDH2*, *ITGA5*, and *ITGB2* are rescued upon co-depletion of *ZEB1* and *ZEB2*, with *ITGA5* gene expression significantly rescued (Fig. 5B). This shows that TRIP12 regulates these EMT cell adhesion genes through suppressing *ZEB1* and *ZEB2* expression. Additionally, *OCLN*, *DSC3*, and *LAMB1* were not rescued upon ZEB1/2 depletion (Fig. 5B). This suggest that there could be ZEB1/2 independent mechanisms which TRIP12 could regulate *OCLN* and *DSC3* as these genes are still rescued with ectopic TRIP12 (Fig. 4A). The expression of LAMB1 is an example of a non-robust target, which also does not decrease robustly in Fig. 4A upon TRIP12 depletion, and acts as a negative control (Fig. 5B).

Next, we performed wound-healing assay to investigate the effect of ZEB1/2 depletion on TRIP12-dependent EMT inhibition. Co-depletion of ZEB1/2 in shTRIP12-2^(6427-6447)^ cells resulted in fewer single cell detachment from bulk cells (Fig. 5C and 5D, Movies S5-S6). Specifically, we see a rescue in cell polarity and cell behavior (Movies S5-S6). Quantitatively, the number of single cells detached from bulk cells is significantly fewer upon ZEB1/2 depletion (Fig. 5D). Overall, this data shows that TRIP12-dependent EMT and metastasis process inhibition is dependent on ZEB1/2 gene expression.

## DISCUSSION

In physiologically normal conditions, TRIP12 targets the tumor suppressor p14ARF for degradation to maintain it at a steady-state level (Chen et al., 2010a). However, this equilibrium changes when there is oncogenic stress (for example, c-Myc amplification) which inhibits degradation of p14ARF by TRIP12, thereby increasing p14ARF levels and inducing OIS (Chen et al., 2013; Chen et al., 2010a). OIS plays a crucial role in preventing cells from turning cancerous and as such is an important player in prohibiting early-stage carcinogenesis (Collado and Serrano, 2006). Based on these findings, changes in TRIP12 level is likely to play an oncogenic role in carcinogenesis and it has been proposed that TRIP12 could serve as a therapeutic target in specific scenarios i.e. when TRIP12 is dysregulated (Collado and Serrano, 2010) or specifically in acute myeloid leukemia where nucleophosmin is attenuated (Chen et al., 2010b). However, our study illustrates a novel role for TRIP12 in suppressing EMT and metastasis-related processes. Specifically, loss of TRIP12 induces an EMT (Figure 3) and results in a higher frequency of single-cell detachment (Fig. 4B and 4C; Movies S1-S4). Conversely, TRIP12 overexpression sensitizes cells to anoikis (Fig. 4D). Additionally, the effects of EMT and single-cell detachment are dependent on ZEB1/2 gene expression (Fig. 5). These results are in line with patient datasets which show a correlation of TRIP12 with distant metastasis-free survival (Fig. 1). These results depict TRIP12’s suppressive role in EMT and suggest an inhibitory role in cancer metastasis.

A plausible reason for the disparity in TRIP12’s function in cancer development between this study and Chen et al (2010a) could be due to the genetic background of the cell model MCF10A used in this study. The immortal MCF10A cell line, which arose spontaneously from mortal mammary epithelial cells, harbors a reciprocal translocation t(3;9) (p14;p21), resulting in the deletion of the *CDKN2A* locus containing p16 and p14ARF (Cowell et al., 2005; Debnath et al., 2003; Soule et al., 1990). The inactivation of the *CDKN2A* locus has also been reported in other immortalized mammary cell lines, illustrating the importance of losing p16 and p14ARF, and in turn loss of OIS, in the immortalization process (Brenner and Aldaz, 1995). While TRIP12 plays a cancer transformation promoting role in p14ARF positive normal fibroblast cells through the degradation of p14ARF resulting in loss of OIS, TRIP12 plays a cancer progression inhibitory role in p14ARF absent immortalized MCF10A cells, which have overcome OIS during the process of immortalization. This represents two opposing functions of TRIP12 in different stages of cancer development, between normal and immortalized stages based on the presence or absence of p14ARF respectively, and adds to the complexity of treatment modalities at different cancer stages. Taken together, it implies that although targeting TRIP12 in early-stage cancer with functional OIS mechanism could be a potential therapeutic approach, targeting TRIP12 in mid to late-stage pre-metastatic invasive breast cancer p14ARF negative patients which do not have a functional OIS may be deleterious.

Although TRIP12 is an E3 ligase involved in protein degradation, this study finds that TRIP12 regulates the expression of multiple cell adhesion genes, mainly through inhibiting ZEB1/2 gene expression. Although the mechanism of how TRIP12 regulates ZEB1/2 expression is not addressed in this study, previous literature has shown that TRIP12 could regulate gene expression through the degradation of epigenetic factors such as the ARID1A complex component BAF57 (Kajiro et al., 2011; Keppler and Archer, 2010) and the Polycomb repressive deubiquitinase (PR-DUB) complex components ASXL1 and BAP1, through a DNA N^6^-Methyladenine sensor network (Kweon et al., 2019). Whether TRIP12 regulates ZEB1/2 through these epigenetic regulators or other regulators requires further investigation.

Apart from cancer, recurrent TRIP12 mutations occurs frequently in patients with neurodevelopmental diseases exhibiting symptoms of autism spectrum disorder, intellectual disability, and craniofacial dysmorphism (Bramswig et al., 2017; O’Roak et al., 2014; Zhang et al., 2017). However, a causal role for TRIP12 in this aspect has not been established. Our finding that TRIP12 regulates EMT could provide some clues on TRIP12’s role in this disease. EMT is an embryonic program required for many processes in embryonic development (Thiery et al., 2009). Specifically, primary EMT during nervous system development drives the formation of migratory neural crest cells, giving rise to peripheral nervous system cells, craniofacial structures, melanocytes and some endocrine cells (Acloque et al., 2009; Thiery et al., 2009). Taken together with our finding, TRIP12 mutations could be affecting EMT during nervous system development, thereby giving rise to neurodevelopmental diseases.

Prognostic markers of metastasis are essential in providing vital information to patients regarding survival and adjuvant treatment eligibility (Weigelt et al., 2005). Here, clinical breast patient datasets show that TRIP12 levels are correlated with low metastasis probability (Fig. 1), suggesting that TRIP12 could potentially be used as a prognostic marker for metastasis prediction and possibly for the stratification of patients for adjuvant treatment. This could both lower metastasis rates in high-risk patients and spare low-risk patients from redundant treatment (Weigelt et al., 2005).

In summary, we have identified a novel EMT and metastasis-related processes inhibitory role for TRIP12 through ZEB1/2 (Fig. 6). In our study, we found several lines of evidence both in the cellular context and in patients’ datasets, which support this newly found function of TRIP12. Clinically, we suggest that therapies to increase TRIP12 levels in mid to late-stage pre-metastatic p14ARF negative breast cancer could be a potential therapeutic approach to ablate metastasis.

**Fig. 6.**
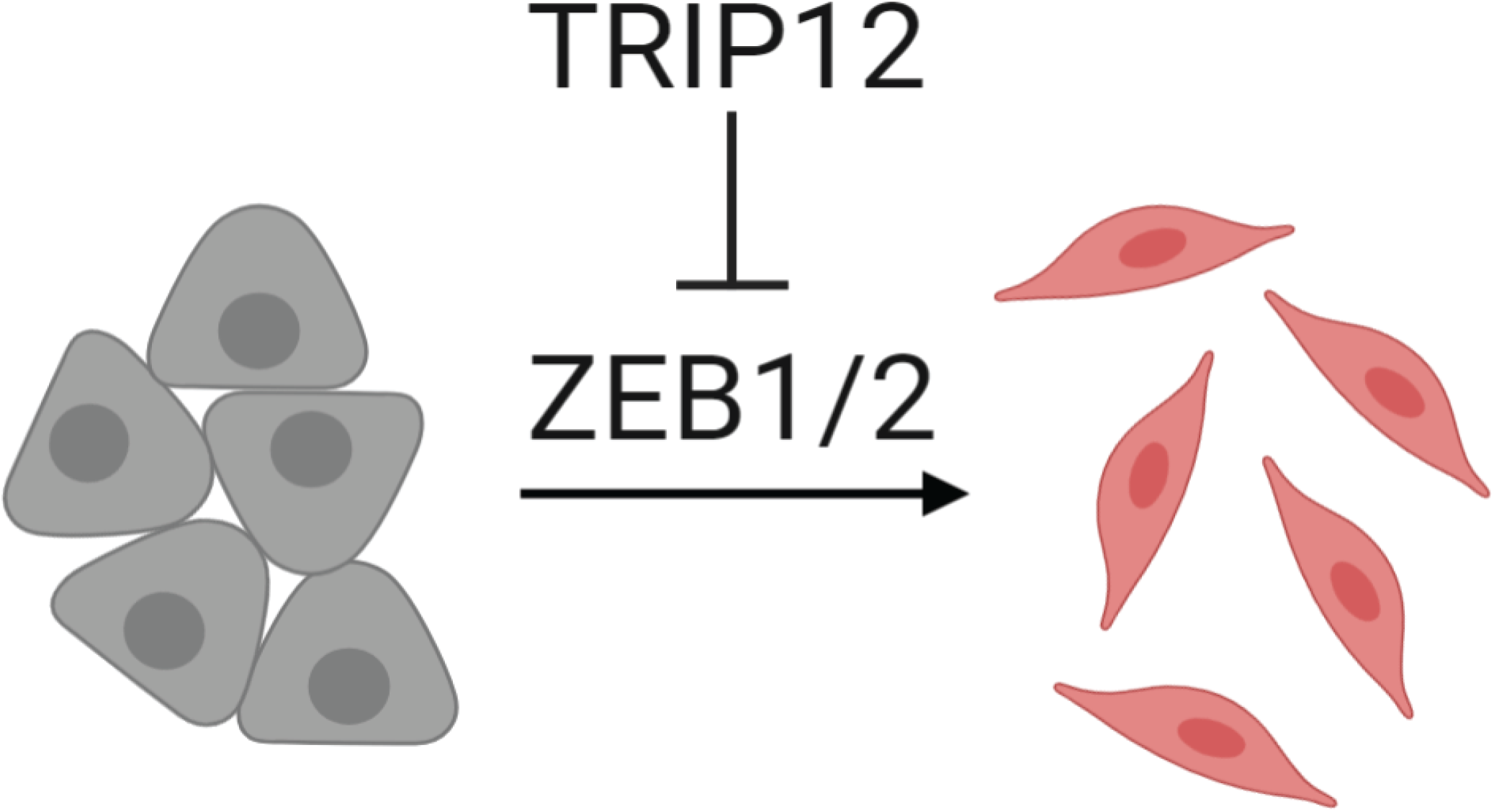
TRIP12 inhibits EMT through repressing ZEB1/2 expression.

## MATERIALS AND METHODS

### Cell culture

MCF10A (ATCC CRL-10317) was grown in Dulbecco’s modified eagle’s media (DMEM)/F12 (1:1) media (Gibco, Cat. No. 11330-032) with 5% horse serum (Gibco, Cat. No. 16050-122) and 1% penicillin-streptomycin (Gibco Cat. No. 15140-122). Additional additives include 20 ng/mL epithelial growth factor (Peprotech, Cat. No. AF-100-15), 0.5 mg/mL hydrocortisone (Sigma-Aldrich, Cat. No. H-0888), 100 ng/mL cholera toxin (Sigma-Aldrich, Cat. No. C-8052) and 10 μg/mL insulin (Sigma-Aldrich, Cat. No. I-1882). HEK293T (ATCC CRL-3216) was grown in DMEM high glucose (Sigma Cat. No. D-5796) with 10% fetal bovine serum (Sigma Cat. No. F-7524) and 1% penicillin-streptomycin (Gibco Cat. No. 15140-122). Cells were grown in a 5% CO_2_ incubator at 37°C. Media was changed 1 day after cells were revived.

### Antibodies

Primary antibodies used were the following: TRIP12 (Bethyl Laboratories, Cat. No. A301-814A), α-Actinin (Santa Cruz, Cat. No. sc-17829), and E-cadherin/CDH1 antibody (BD Biosciences, Cat. No. 610182).

### shRNA

shRNA was cloned into pLKO.1 vector using restriction enzyme sites *Age*I (NEB, Cat. No. R3552) and *Eco*RI (NEB, Cat. No. R3103). pLKO.1 puro was a gift from Bob Weinberg (Addgene plasmid # 8453; http://n2t.net/addgene:8453; RRID: Addgene_8453). shRNA sequences used in this study are as follows:

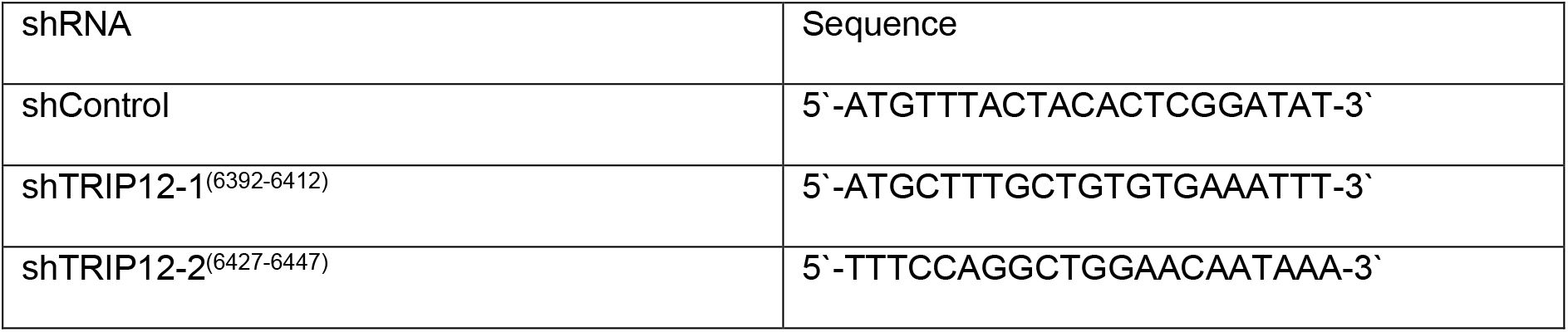

### siRNA

siRNA used in this study are as follows:

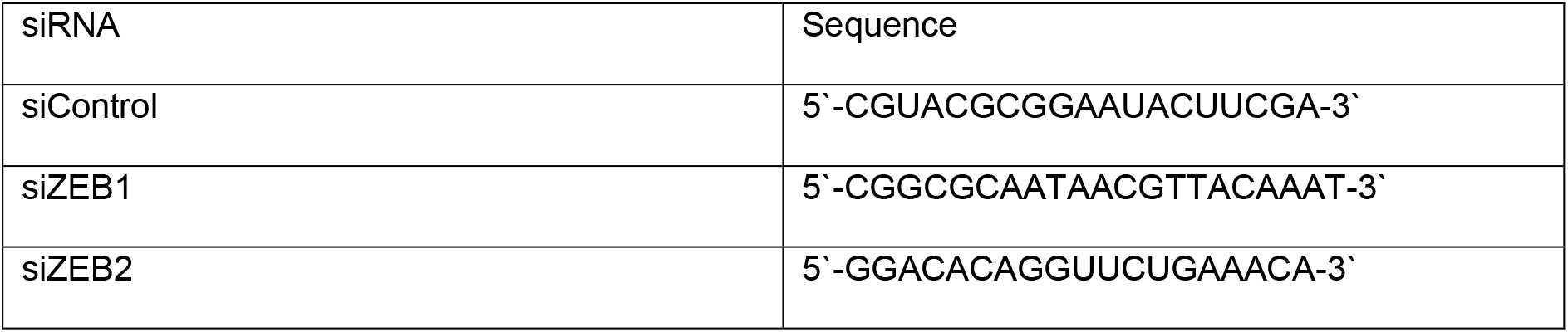

### siRNA transfection

siRNA was transfected using Lipofectamine RNAimax (Thermo Scientific, Cat. No. 13778150) according to the manufacturer’s protocol. Transfection mix was prepared by addition of 1 mL of Opti-MEM (Thermo Scientific, Cat. No. 31985070), 5 µL of Lipofectamine RNAimax and 20 nM total siRNA (for RNA analysis) or 10 nM total siRNA (for cell migration/ wound-healing assay) required for a total volume of 5 mL. The mixture was mixed and left standing for 30 min before addition to 8×10^5^ cells in a 10 cm dish with total volume of 5 mL (complete media with cells and transfection mix). After 6 h, fresh media was used to replace the transfection media. Cells were harvested after 48 h from time of transfection.

### Plasmid

The pAcGFP-C1-TRIP12 plasmid was provided by Professor Jiri Lukas. TRIP12 CDS was subcloned into the LHCX vector (Clontech, Cat. No. 631511) using a single *Not*I site introduced into the LHCX vector through the *Hind*III site.

### Generation of stable cell lines

Virus was generated by transfecting 3×10^6^ HEK293T cells with plasmids using Lipofectamine 2000 (Thermo Scientific, Cat. No. 11668019) according to the manufacturer’s protocol. Viruses were harvested, filtered through 0.45 μm filter, and used to infect 5×10^5^ MCF10A cells (seeded 1 day before) with polybrene (Sigma-Aldrich, Cat. No. S2667) (8 μg/mL). After 24 h, the media was replaced with fresh media and left for 24 h for cell recovery. Media containing puromycin (Sigma, Cat. No. P9620) (1 μg/mL) or hygromycin (Invitrogen, Cat. No. 10687010) (50 μg/mL) was added and changed every 48 h till mock-transfected cells died. Cells were grown in antibiotics containing media for 2 weeks for the creation of stable cells.

### RNA extraction

RNA was extracted using TRIZOL reagent (Life Technologies, Cat No. 15596026). One milliliter of TRIZOL was added to harvested cells and mixed by pipetting up and down several times. To ensure efficient lysis, the mixture was vortexed for 15 sec. The mixture was left at room temperature for 5 min to allow efficient dissociation of nucleic acid-protein complexes. Chloroform (Sigma, Cat. No. C2432) equivalent to 20% of the amount of TRIZOL used (200 μL) was added, vortexed for 15 sec and left to settle for 3 min. The mixture was centrifuged at 12,000×g for 15 min. The top layer after centrifugation, equivalent to ~50% of the volume of TRIZOL used (500 μL), was transferred to another tube. Volume equivalent to 50% of TRIZOL used (500 μL) of isopropanol (Sigma, Cat. No. I9516) was added, mixed and left at room temperature for 10 min. The mixture was centrifuged at 12,000×g for 10 min. The supernatant was removed, and the pellet washed with 1 mL of 75% ethanol. Centrifugation was done at 7,500×g for 5 min. The ethanol wash step was repeated, and the pellet was left to dry for 5 min on a benchtop. The RNA pellet was dissolved in RNase-free water (Santa Cruz, Cat. No. sc-204391).

### RT-qPCR

cDNA was synthesized using the iScript cDNA synthesis kit (Bio-Rad, Cat. No. 170-8891) following the manufacturer’s protocols. qPCR was performed using iTaq Universal SYBR Green Supermix (Bio-Rad, Cat. No. 172-5125) on a ThermoFisher Scientific QuantStudio 3 or 5 Real-Time PCR system. CT values were normalized to actin and to control cells to achieve ΔΔCT values and expressed as relative expression. The primers used for qPCR are as follows:

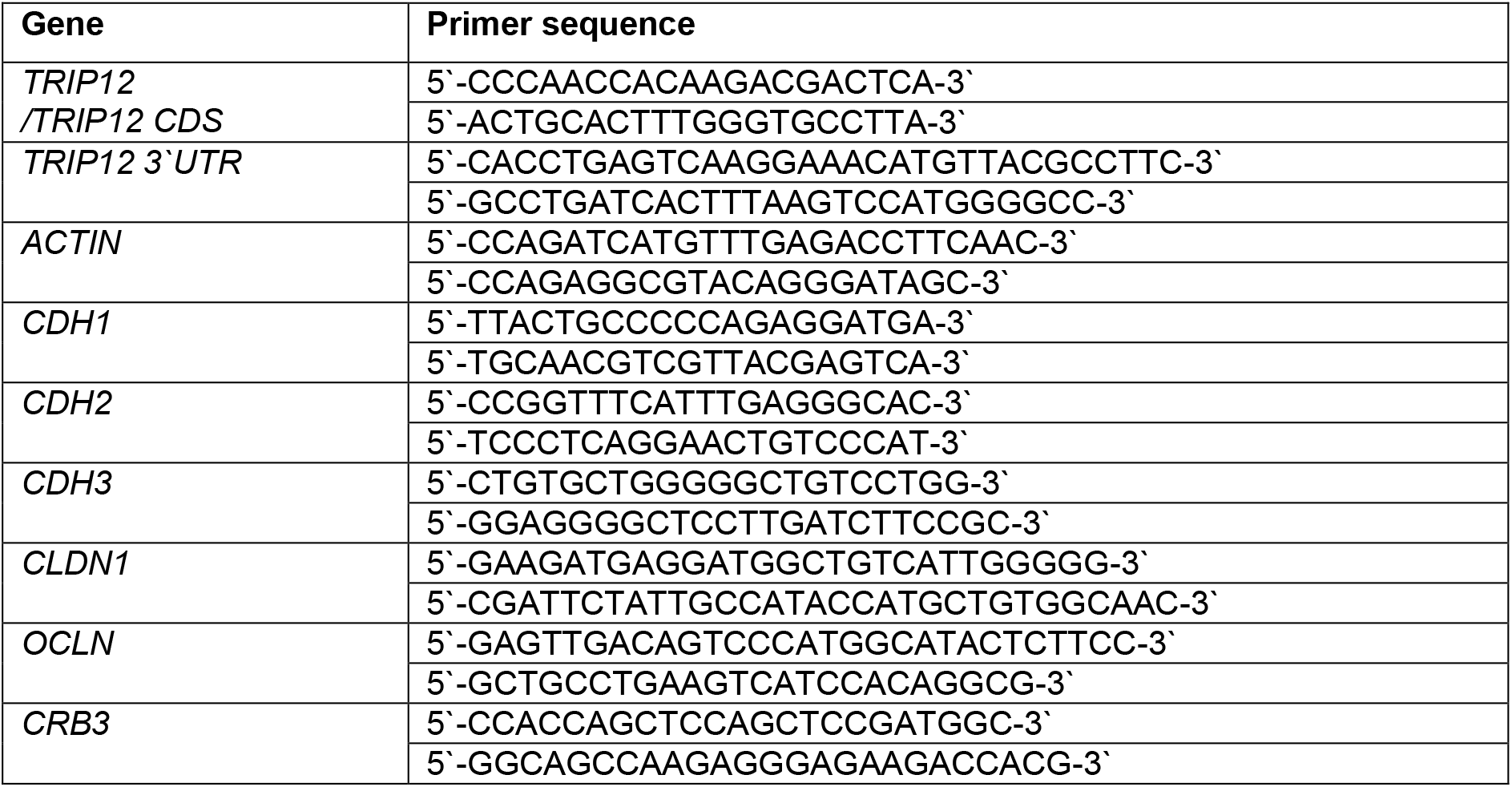
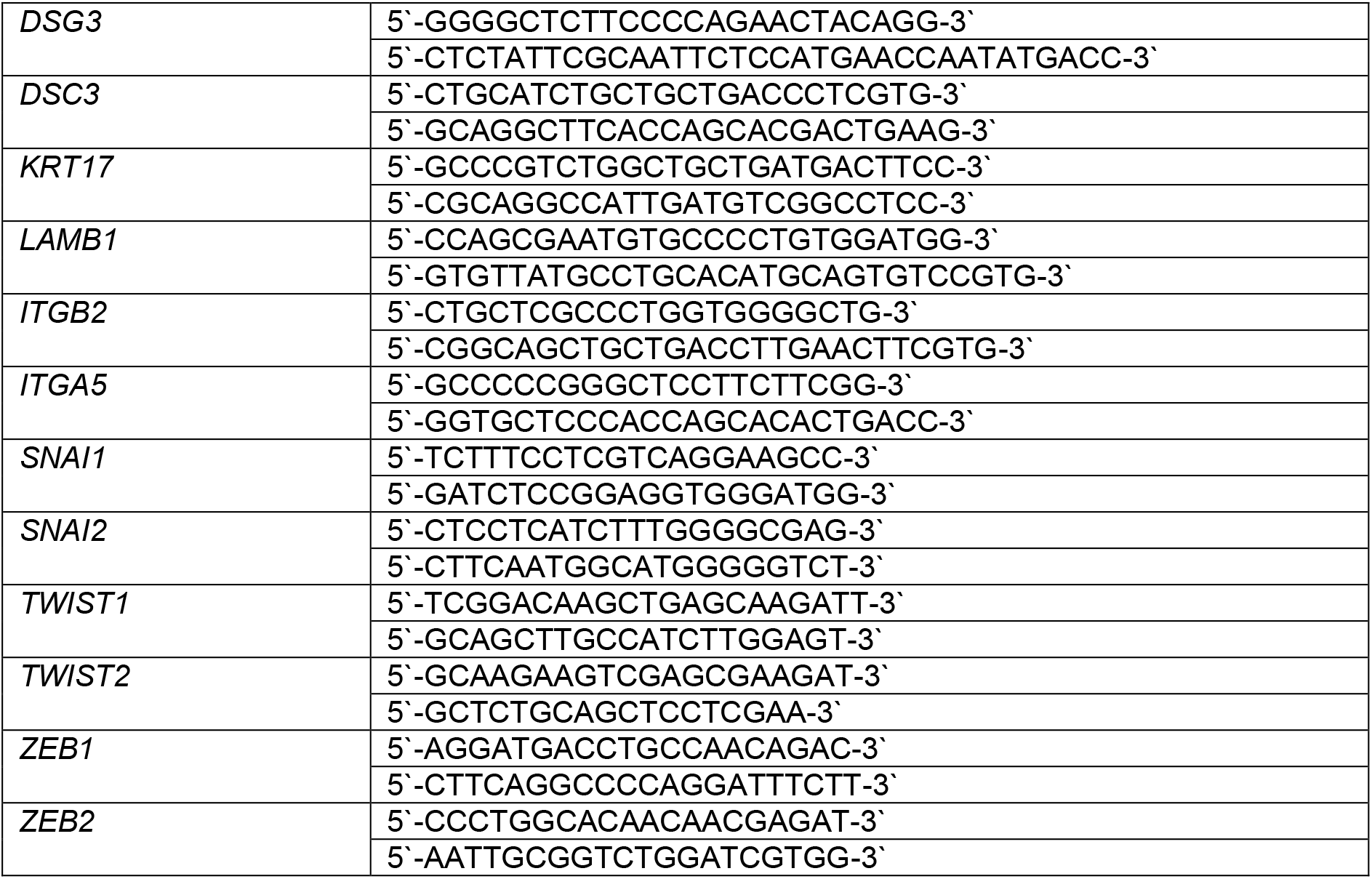

### Cell migration/ wound-healing assay

Cells were seeded at low confluency (3×10^5^ cells in a 10-cm plate) for at least 2 passages with newly constituted media before seeding for cell migration/wound-healing assay. Cells were seeded in triplicates at 4×10^5^ cells per well in a 24-well plate in serum-free media with additives overnight. A wound was created using a 10 μL pipette tip by gently scratching the well from top to bottom in a vertical motion. Cells were washed four times with 0.5 mL PBS to remove detached cells and 2 mL of serum-free media with additives was added after the last wash. Live imaging of cells was performed using Zeiss Axio Z1 with a cell observer (Zeiss) with image capture every 0.5 h until wound closure. Quantification was performed using “ImageJ”. Polygon selection (“ImageJ”) was used to trace the boundary of bulk cells at a specified timepoint to measure the remaining wound area. Quantification of single-cells was performed by counting the number of single-cells in images when the remaining wound area is 35% - 55% of the total image area.

### Anoikis assay

Fifty microliters of 1 % agarose was added into each well of a 96-well plate and left to solidify. Cells were counted and 2×10^4^ cells in 100 μL were seeded per well in triplicates per condition. At timepoints 0 h and 24 h, 20 μL of CellTiter 96 Aqueous One Solution cell proliferation assay (MTS) (Promega, Cat. No. G3581) was added to each well. The plate was incubated at 37°C and the absorbance of each well was taken at 490 nm at each hour for a total of 3 h. The background absorbance in each well was normalized by subtracting the absorbance value of a well without cells to increase the signal-to-noise ratio. Twenty-four hours reading was normalized to 0 h reading and expressed as % viability.

### Immunofluorescence staining and imaging

Cells were seeded at 3×10^5^ cells per well in a 6-well plate and grown in a 5% CO_2_ incubator at 37°C for 24 h. Cells were washed once with PBS and fixed in 3.7% (w/v) paraformaldehyde (Sigma, Cat. No. P6148) for 30 min. Following which, cells were washed thrice with PBS, permeabilized with 0.1% Triton-X (Naclai Tesque, Cat. No. 25987-85) for 10 min, washed thrice with PBS, blocked with 3% BSA containing 0.1M glycine for 30 min, washed thrice with PBS, and incubated with primary antibody (1:500 in 3% BSA dissolved in PBS) for 1.5 h at room temperature or overnight at 4°C. Fixed cells were washed thrice with PBS and incubated in Alexa Fluor 488 Goat anti-mouse IgG secondary antibody (Thermo Fisher, Cat. No. A-32723) (1:500 in 3% BSA dissolved in PBS) for 30 min at 37°C. Cells were washed five times with Milli-Q water and stained with Hoechst 33342 (Santa Cruz, Cat. No. SC-200908) dye at 1 μg/mL in Mili-Q water for 30 min. Images were taken using a Zeis Axiovert.A1 microscope (Zeis).

### Statistical analysis

The student t-test was used for significance testing. p-values are depicted as such (*, *p* < 0.05; **, *p* < 0.01; ***, *p* < 0.001). Where *p* > 0.05, the p-value is shown to 3 decimal points. The number of independent replicates used for statistical analysis in all cases is indicated in the respective figure legends. N represents biological replicates. Error bars were plotted using either standard deviation (SD) or standard error of the mean (SEM) and are indicated in the respective figure legends.

## ACKNOWLEDGMENTS

We thank the members of the Jha laboratory for helpful discussions and comments. We thank Professor Jiri Lukas for the pAcGFP-TRIP12 plasmid used in this study (Gudjonsson et al., 2012).

## AUTHORS’ CONTRIBUTIONS

Conception and design: K.K. Lee, S. Jha

Development of methodology: K. K. Lee, D. Rajagopalan, S. Jha

Acquisition of data: K. K. Lee, D. Rajagopalan

Analysis and interpretation of data: S. S. Bhatia, K.K. Lee, D. Rajagopalan, S. Jha

Writing, review and/or revision of the manuscript: K.K. Lee, D. Rajagopalan, S. S. Bhatia, S. Jha, W. J. Chng

Administrative, technical, or material support: K.K. Lee, S. Jha

Study supervision: S. Jha, W. J. Chng

